# The Dry Truth: Hair-Wetting Improves Dry Electrode EEG Signal Quality

**DOI:** 10.64898/2026.09.21.753100

**Authors:** Yingqi Huang, Abele Michela, Victor Férat, Cristina Colangelo, Kishen Senziani, Laurence Castonguay, Serge Vulliémoz, Tomas Ros

**Affiliations:** CIBM Center for Biomedical Imaging, Geneva, Switzerland; Department of Basic Neurosciences, University of Geneva, Geneva, Switzerland; Department of Psychology, University of Montreal, Montreal, Canada; Department of Clinical Neurosciences, University of Geneva, Geneva, Switzerland; Swiss Police Institute, Neuchâtel, Switzerland

**Keywords:** EEG, dry electrodes, signal quality, BCI, Resting State

## Abstract

With the rapid advancement of clinical neuroscience and Brain-Computer Interfaces (BCIs), there is an increasing demand for convenient, user-friendly EEG recording methods suitable for diverse environments, including mobile and home-based settings. Traditional gel-based EEG systems, while reliable, are inconvenient and time-consuming, whereas dry electrode systems tend to suffer from elevated noise levels.

In this study, we investigated a novel methodological manipulation aimed at enhancing dry electrode signal quality: wetting the hair directly with tap water to improve scalp electrode conductivity. To this end, we recruited 22 healthy participants and compared their resting-state (RS) EEG activity across three experimental conditions (dry, semi-dry, and gel) within a single-session design. Specifically, we analyzed electrode impedance, spectral power (SP), and functional connectivity (FC).

Our results show that basic hair-wetting substantially improved mean electrode impedance by 75.7% relative to the dry condition, reduced the proportion of bad channels from approximately 40% to 15%, and increased spectral-power similarity with gel recordings by 7.7% during eyes-closed recordings, and improved FC similarity by approximately 56-70% across frequency bands.

Although this method does not fully match the signal quality of traditional gel-based systems, it represents a promising compromise, enhancing EEG data quality under suboptimal conditions. This approach offers a practical and non-invasive means to improve EEG signal quality, ultimately expanding real-world applications of EEG.

## Introduction

Due to its high temporal resolution, portability, and relatively low cost, electroencephalography (EEG) has become a fundamental tool in both clinical neuroscience and research. Clinically, EEG is widely used for diagnosing and monitoring neurological disorders such as epilepsy, sleep disorders, traumatic brain injury, and encephalopathies (Silva & Niedermeyer, 2012). In recent years, its role has expanded into neuropsychiatry, where EEG biomarkers are being investigated for conditions like depression, schizophrenia and attention-deficit/hyperactivity disorder (ADHD) (Michel & Murray, 2012; Newson & Thiagarajan, 2019).

Beyond diagnosis, the high temporal resolution of EEG is advantageous for developing brain-computer interfaces (BCIs) that convert real-time neural activity into “neurofeedback” control signals (Nicolas-Alonso & Gomez-Gil, 2012). These systems serve various purposes, from motor rehabilitation to cognitive training and affective self-regulation. EEG-based BCIs are being used not only as therapeutic tools for clinical populations but also by healthy individuals seeking to enhance mental wellness, manage stress, and boost cognitive performance (Gruzelier, 2014; Voigt et al., 2024). The growth of mobile platforms and consumer-grade EEG devices reflects a shift toward brain monitoring outside typical clinical settings (Enriquez-Geppert et al., 2019). Recent advancements in tele-neurotechnology and the decentralisation of mental health treatments have led to the creation of tools for patients to use EEG at home over several months with or without clinical supervision (Biondi et al., 2024; Micoulaud-Franchi et al., 2015; Ölçüoğlu, 2025).

Nevertheless, success in these areas relies on user-friendly EEG systems that can capture reliable brain data in everyday environments (Niso et al., 2023). Despite its mobility, the widespread use of EEG outside controlled settings is limited by practical challenges, especially those related to comfort, self-application, and signal quality (Abiri et al., 2019; Radüntz & Meffert, 2019). Gel-based EEG systems provide high signal quality and low impedance but take time to set up, need trained staff, and can be uncomfortable for long periods, making them unsuitable for home use (Ng et al., 2022). In contrast, dry electrode systems promise quicker setup and more convenience, but often have high impedance and much lower signal reliability (Chi et al., 2012; Mathewson et al., 2017; Sendi, 2025). A significant drawback of current dry electrode systems is the frequent occurrence of poor quality signal or “bad” electrodes, which can affect signal quality and the accuracy of analyses, as well as lead to more data rejection (Lewis et al., 2024; Tăuţan et al., 2013). This decline in signal quality impacts not only visual analysis of EEG time-series (e.g., in epilepsy) but also its quantitative biomarkers, such as power spectral density and functional connectivity, both crucial in clinical and BCI applications (Moumane et al., 2024; Ng et al., 2022). Despite enhancements in materials and design, the ongoing presence of “noisy signals” in dry EEG is a major hurdle for using these systems in unsupervised or home environments where expert signal quality checks and technician adjustments are not feasible (Barbey et al., 2022).

This study explores a simple, low-cost method for enhancing dry-electrode performance: wetting the hair with water before EEG cap application. We hypothesised that this relatively simple manipulation may improve electrode-scalp conductivity by reducing hair volume (Mathewson et al., 2017) and/or increasing electrode-scalp surface contact area. However, this approach has not been systematically evaluated to date, highlighting the need for its empirical investigation.

In this study, we specifically compare the resting state recording based on three electrode preparation methods within a single-session design (i.e. one sitting within the same subject): dry, semi-dry (i.e. dry with wet hair), and gel-based (Kleeva et al., 2024). We then assessed signal quality through direct comparisons of electrode impedance, power spectral density (PSD), and functional connectivity (FC) measures. Impedance provided an indication of electrode–scalp contact, while PSD allowed us to examine whether the spectral characteristics of dry-electrode recordings resembled those of the gel-based reference. Finally, we assessed FC using phase locking. This measure complements the PSD analysis by evaluating the consistency of inter-electrode phase relationships rather than spectral power alone. Together, these measures allowed us to evaluate both signal quality and the similarity of key EEG features. By testing a simple way to improve dry-electrode recordings, this study may inform the development of more practical EEG systems for home-based monitoring and BCI applications (Sendi, 2025).

## Materials and methods

### Participants

A total of 22 healthy adult participants (age range: 18–30 years, 6 males, 16 females) were recruited for this study. All participants provided informed consent in accordance with the Declaration of Helsinki, and the study was approved by the institutional ethics committee.

### Experimental Conditions

Each participant underwent resting-state EEG recordings (Figure 1a) under three different electrode preparation conditions in the following order:

**Figure 1.**
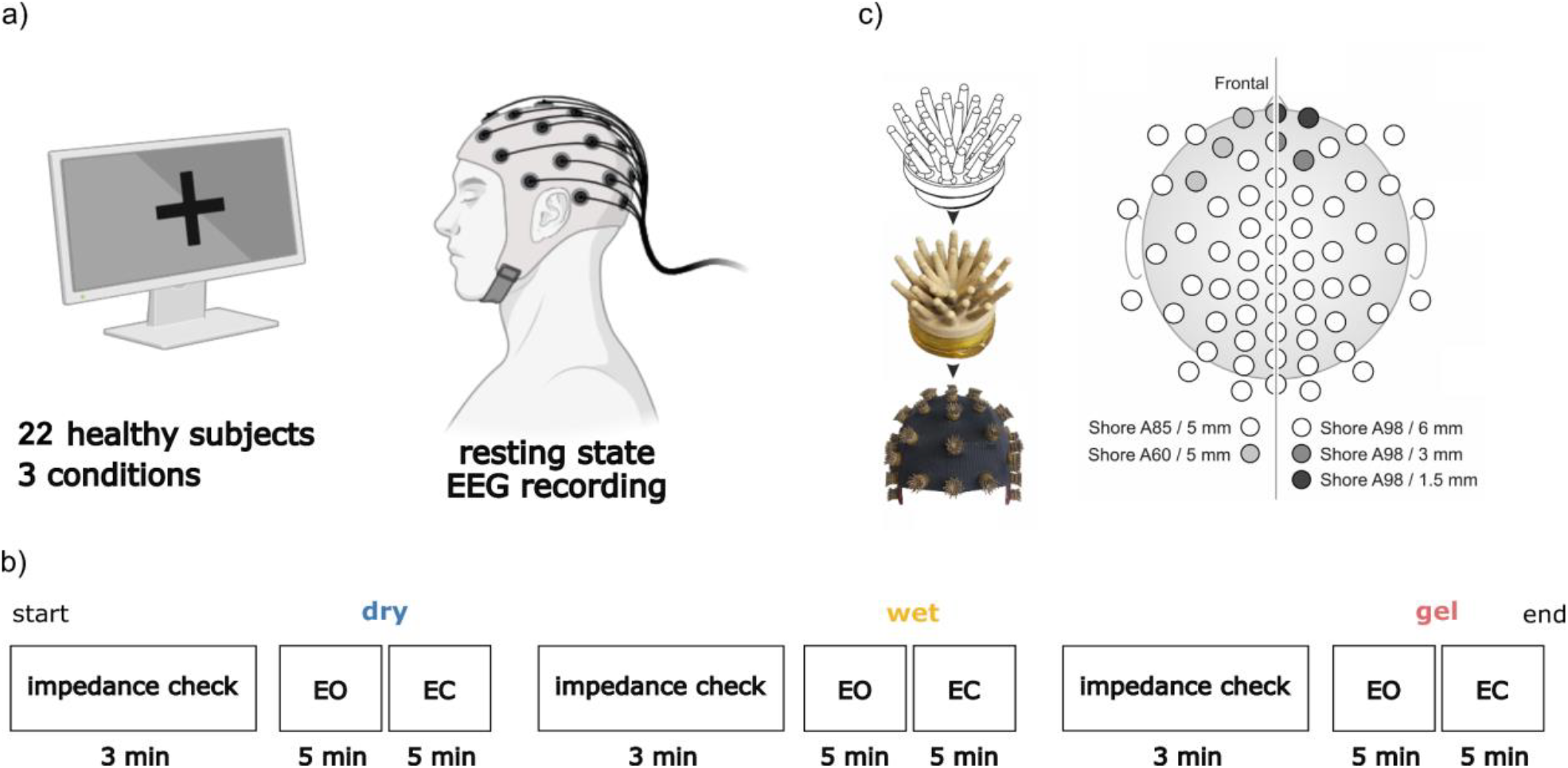
Experimental protocol. (a) 22 healthy participants are recruited for resting state EEG recording. (b) The experimental protocol. (c) The flower electrode schematic and an example of a 64-channel dry cap montage (Warsito et al., 2023).

1. **Dry**: The cap was applied as-is with no additional preparation.
2. **Semi-dry / “wet”**: Participants’ hair was completely wetted using tap water before dry-cap application, without applying conductive gel. We hereafter use the term ‘wet’ to refer to this semi-dry condition.
3. **Gel**: Following drying of hair with a hair dryer, standard EEG gel was applied to each electrode. This served as the gold-standard reference condition.

As shown in Figure 1b, each condition’s resting-state recording consisted of 5 minutes with eyes open and 5 minutes with eyes closed. Participants were instructed to remain relaxed, minimise movement, and avoid excessive blinking throughout the recordings. Conditions were presented in a fixed order: dry, wet, and gel. An impedance check was performed at the beginning of each session. Based on the impedance values, the researchers adjusted the cap electrodes. Impedance values were recorded after adjustment.

### EEG Recording Setup

EEG for the dry and wet conditions was recorded using a 32-channel dry electrode cap equipped with specialised “flower”-shaped (Figure 1c) Ag/AgCl-coated electrodes (Waveguard touch, ANT neuro, Netherlands) designed to part the hair and maintain contact with the scalp without the need for conductive gel (Warsito et al., 2023). This setup utilised an ear-linked reference. For the gel condition, the conventional gel-based cap was equipped with 64 electrodes (Waveguard original, ANT neuro, Netherlands), with 29 electrodes overlapping with the dry cap montage. Data were originally sampled at 500 Hz. Impedance values for each channel were recorded prior to each session, during the impedance check, using the manufacturer’s built-in impedance check feature.

### Signal Preprocessing

For comparison across EEG systems, data from the 29 electrodes common to all systems were extracted. EEG data were preprocessed using the MATLAB-based automated denoising pipeline HAPPE (Gabard-Durnam et al., 2018), which includes several standardized denoising procedures. The data were first downsampled to 250 Hz, followed by band-pass filtering between 0.5 – 45 Hz using an FIR filter *pop_eegfiltnew* as an implemented function in EEGLAB. Bad channels were identified and removed based on two criteria: impedance levels and the normalized joint probability of the average log power. Channels exceeding three standard deviations from the group mean were classified as bad and excluded. Independent component analysis (ICA) was then applied, and the identified bad channels were interpolated post-decomposition. Subsequently, ICA components identified as artifacts were automatically rejected.

### Power Spectral Density

PSD was computed using Welch’s method, where absolute power and relative power were then analyzed across conventional EEG frequency bands: delta (1–4 Hz), theta (4–8 Hz), alpha (8–12 Hz), beta (12–30 Hz), gamma (30–45 Hz) and across broadband. To evaluate the data quality, the correlations of relative spectral power of paired subjects and paired channels are compared (Kleeva et al., 2024; Moumane et al., 2024). After concatenating each channel’s EEG signal of each subject, the cross-signal correlation is evaluated by Pearson’s value (Tautan et al., 2014).

### Functional connectivity (FC)

To assess functional connectivity, Phase Locking Value (PLV) was computed between electrode pairs for each condition. The PLV quantifies the consistency of phase differences between two signals across time (Lachaux et al., 1999), providing a measure of synchronisation (Arnulfo et al., 2015), which makes it less sensitive to amplitude fluctuations and noise. We then calculated the spatial correlation of the PLV matrices (i.e. their upper triangular entries) between different conditions to estimate their similarity (Sareen et al., 2021).

### Statistics

In this study, non-parametric statistical analyses were performed due to the non-Gaussian distribution of the data. Multiple-group comparisons in unpaired measurements were assessed using the global Kruskal–Wallis test, and for the post hoc pairwise comparison between groups, we used the Mann–Whitney U test for unpaired features. Whereas paired (i.e. within-subject) features between different conditions were evaluated using the Friedman test for multiple-group comparisons, and using Wilcoxon signed-rank tests for the post hoc pairwise comparisons between groups. All statistical tests were two-tailed with an alpha threshold of 0.05. When applicable, post-hoc pairwise comparisons were performed with Bonferroni correction for multiple groups testing, using the MATLAB function *multcompare()*. Significance levels are indicated as: *\* 0*.*05 > p ≥ 0*.*01; ** 0*.*01 > p ≥ 0*.*001; *** 0*.*001 > p*.

## Results

### Impedance check and pipeline performance

Average impedance values were computed for each condition and compared statistically. As illustrated in Figure 2a, all pairwise comparisons between impedance levels were significant (Kruskal-Wallis H(2) = 207.9; wet vs. dry: d = -1.32, corrected p < 0.001; wet vs. gel: d = 1.83, corrected p < 0.001). Notably, the dry cap condition exhibited the highest impedance (mean: 350, SD: 280 kΩ), followed by the wet cap (mean: 85, SD: 58 kΩ), and the gel cap condition, which demonstrated the lowest impedance (mean: 1.7, SD: 4.2 kΩ).

**Figure 2.**
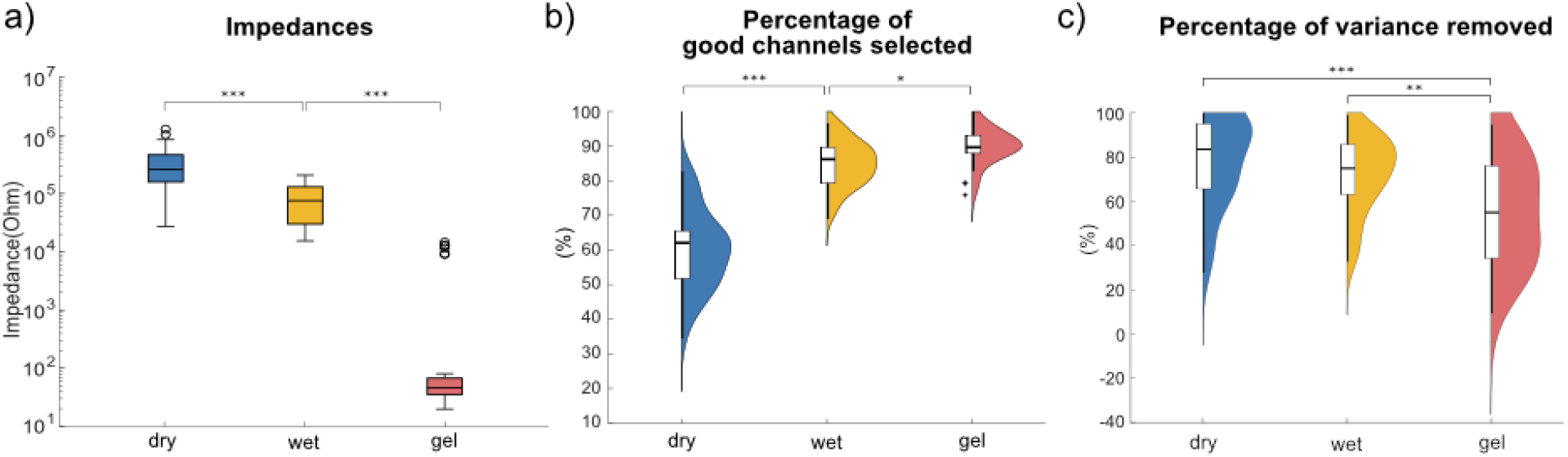
EEG Signal Quality Metrics. (a) Impedance recorded before the EEG task. (b) Percentage of good channels selected by HAPPE. (c) Percentage of variance removed.

According to the data quality check report by HAPPE, as depicted in Figures 2b and 2c, dry electrodes showed the poorest performance, with approximately 40% of channels classified as “bad”, defined as channels exhibiting high impedance or displacement and whose normalized joint probability of average log power in the 1–125 Hz range deviated by more than three standard deviations from the mean (Gabard-Durnam et al., 2018). In contrast, wetting the hair before applying dry electrodes substantially improved signal quality, reducing the proportion of bad channels to around 15%. Gel-based electrodes remained the gold standard, with only about 10% bad channels (Friedman test, chi(2) = 71.51, wet vs. dry: d = 2.63, p<0.001 corrected; wet vs gel: d = 1.69, p = 0.030 corrected). The variances removed between three conditions were also compared with Friedman test, chi(2) = 24.59 ( wet vs dry: d = -0.31, p= 0.21 corrected; wet vs gel: d = 0.81, p= 0.006 corrected).

These results show that the dry condition consistently yielded lower signal quality, as indicated by a high electrodes’ impedance, low percentage of good channels selected by HAPPE and a higher proportion of variance attributed to noise components. While the wet cap condition showed a significantly lower electrodes’ impedance, higher percentage of good channels compared to the dry cap, indicating a better data quality (Lewis et al., 2024), it still did not match the gel cap condition, which achieved the best retention of the data.

### Spectral Power

Figure 3a-d shows the mean absolute and relative PSD across all the participants with their confidence intervals, for the three electrode preparation conditions during eyes open and eyesclosed RS recordings. Overall, these plots show the PSD profiles obtained with wet and gel conditions showed a high degree of similarity (EC: r = 0.7 ± 0.19; EO : r = 0.79 ± 0.14). In the eyes-closed condition, a clear alpha peak is clearly visible, particularly for the wet and gel cap recordings. Compared to the dry condition, where the alpha peak is not as evident, wetting the hair can significantly improve the PSD profile.

**Figure 3.**
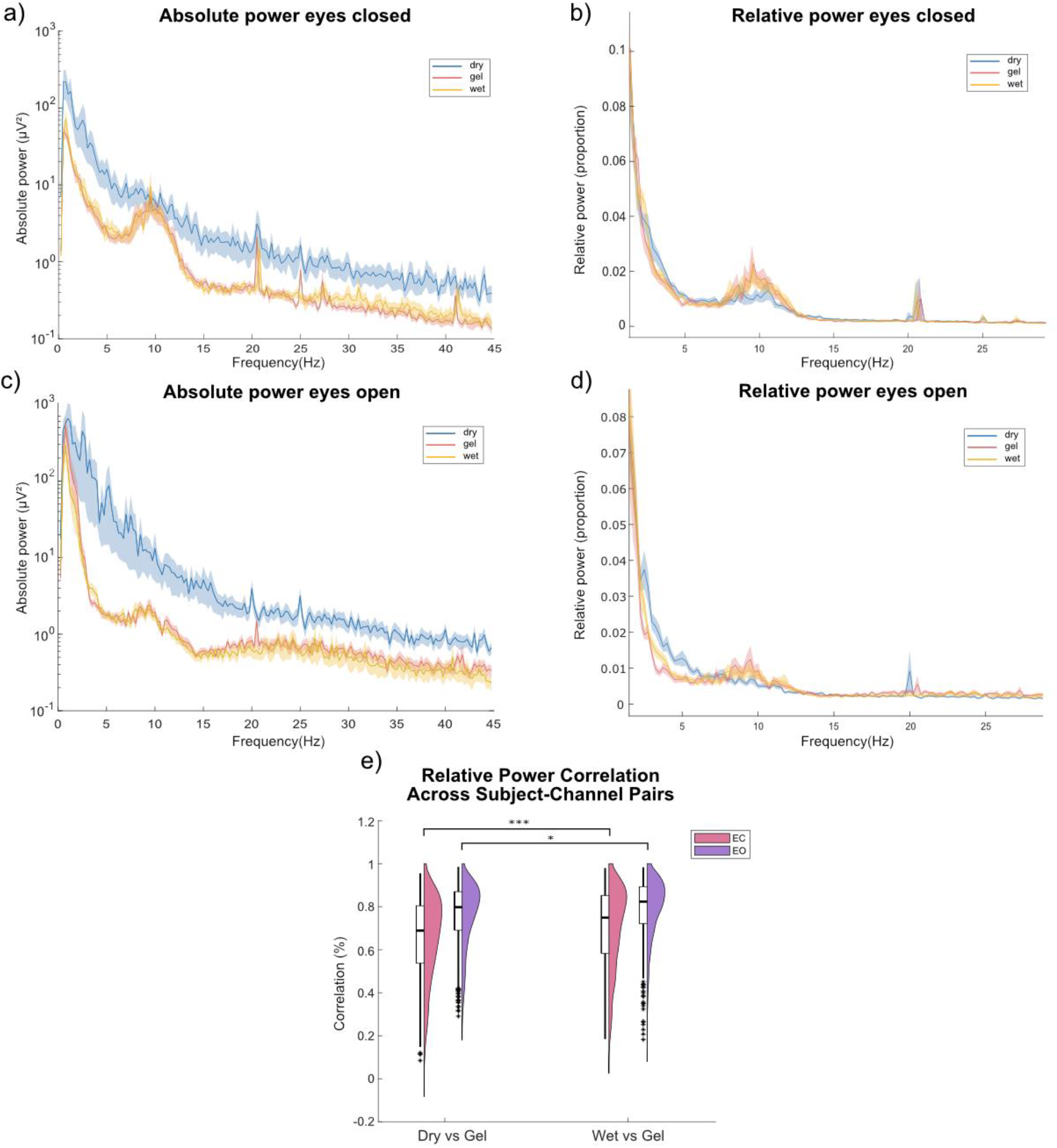
Power Spectral Density. Power of three conditions with their confidence interval. (a/c) Eyes closed/eyes open absolute power from 0.5 to 45 Hz. (b/d) Eyes closed/eyes open relative power from 0.5 to 30 Hz. (e) Intra-subject-channel pairs relative PSD power correlation across broadband.

As demonstrated in Figure 3(e), PSD correlations between conditions were computed for each matched subject–channel pair. With eyes closed, the correlation between wet-gel (mean: 0.70, SD:0.19) is dominant compared to dry-gel (mean: 0.65, SD:0.19). With eyes open, the global correlation increased and the difference between conditions was smaller, with dry-gel correlations of mean: 0.76, SD: 0.14, and wet-gel correlations of mean: 0.79, SD: 0.14. The Wilcoxon test reveals significant differences between dry-gel and wet-gel correlations in both eyes closed recordings (Z = -6.33, dz = -0.25, p < 0.001) and eyes open recordings (Z = -3.48, dz = -0.16, p < 0.001), confirming that electrode preparation had a significant effect on signal quality.

To further explore whether the distributions of the relative power over the scalp are similar between different conditions, the topoplots were drawn for each frequency band. As shown in Figure 4, compared to dry-condition topoplots, wetting the hair before applying dry electrodes restores topographical specificity and has fewer regions of interest that are significantly different from gel, especially in the theta and alpha frequency bands. Gel-based electrodes remained the gold standard, consistent topographical specificity across all frequency bands.

**Figure 4.**
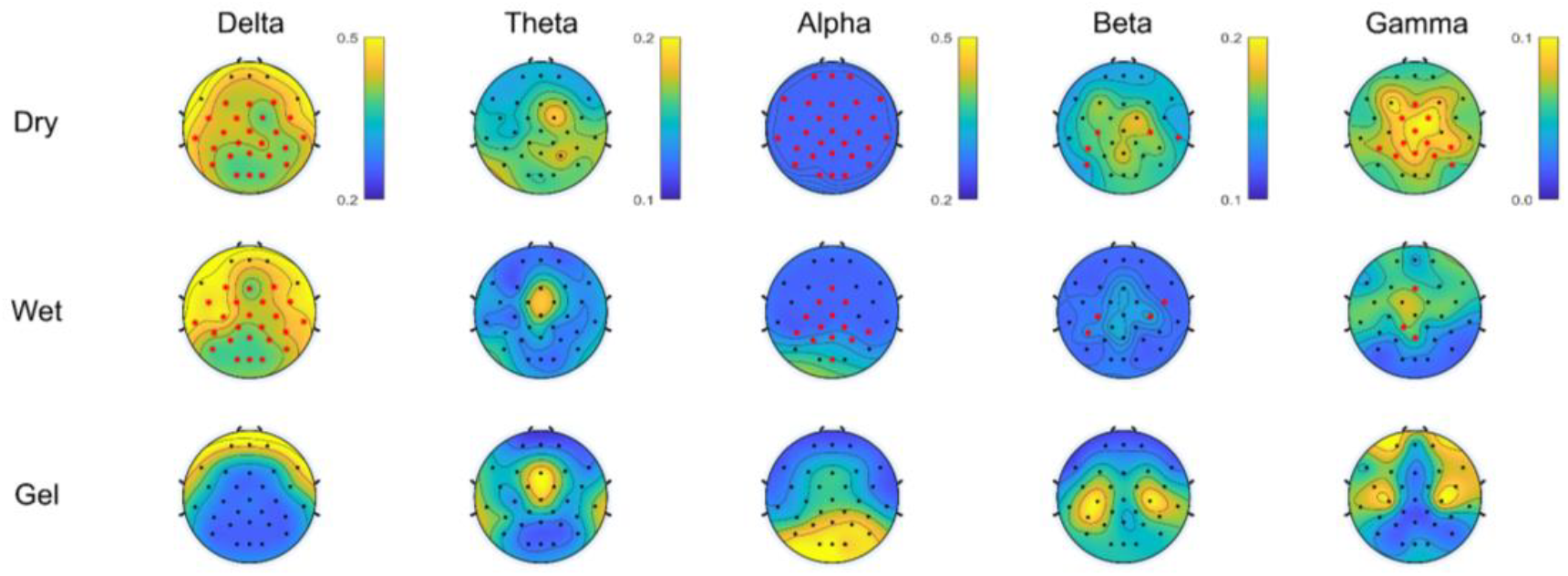
Relative power topoplots of 3 conditions for each frequency band. Asterisks indicate significant differences with the gel condition (p<0.05).

### Functional Connectivity

As indicated by the red line in Figure 5(a), intra-subject similarity of functional connectivity was quantified by correlating PLV values across subject–channel pairs for each frequency band (Figure 5(b)). The results show that wet and gel always have a significantly higher similarity, compared to dry data in all five frequency band (*Delta: Wilcoxon test, Z = -3*.*59, d = -1*.*02, p < 0*.*001; theta: Wilcoxon test, Z = -3*.*85, d = -1*.*28, p < 0*.*001; alpha: Wilcoxon test, Z = -3*.*8, d = -1*.*29, p < 0*.*001; beta: Wilcoxon test, Z = -3*.*94, d = -1*.*35, p < 0*.*001; gamma: Wilcoxon test, Z =-3*.*88, d = -1*.*69, p < 0*.*001*). Wet-gel FC similarity exceeded dry-gel similarity across all frequency bands, with absolute increases ranging from +0.18 to +0.24 in correlation values. Based on group means, this corresponded to relative increases of 55.7% in delta, 57.6% in theta, 60.7% in alpha, 58.6% in beta, and 70.2% in gamma (Table S1). This analysis revealed that PLV connectivity patterns in the wet condition were more reproducible and systematically aligned with the gel condition than those observed in the dry condition.

**Figure 5.**
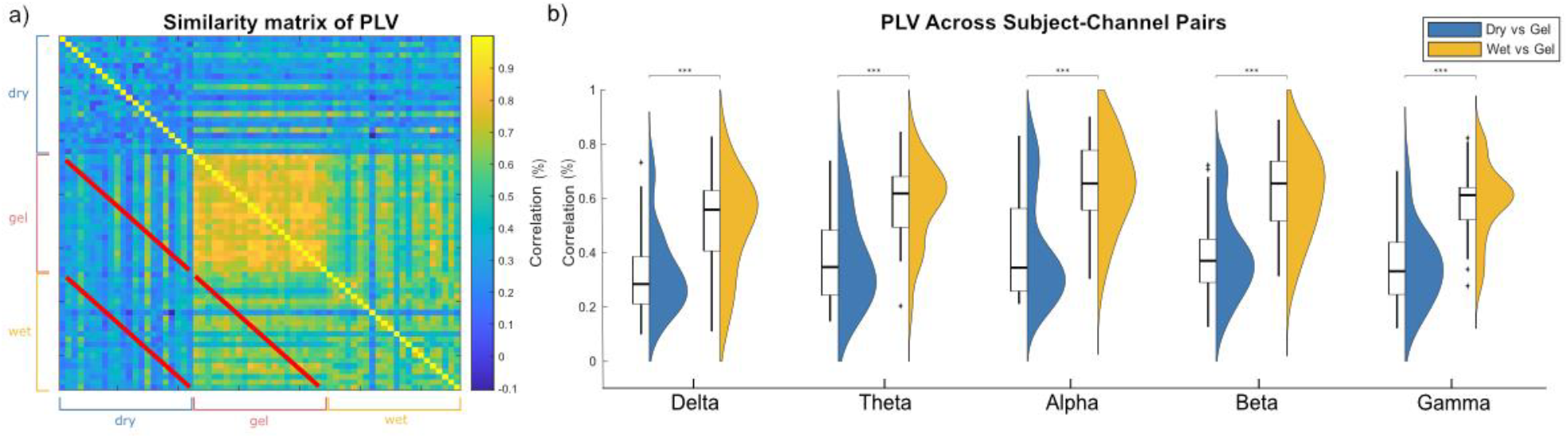
(a) Phase-Locking Value (PLV) similarity matrix across all subjects with eyes closed in all three conditions (dry, gel and wet). This PLV matrix was calculated from broadband (0.5 - 45 Hz) EEG. The red line indicates the pairwise PLV correlations, where we calculated intra-subject PLV correlation between different conditions. In the second subplot this feature was compared in different frequency bands. (b) The violin plots of intra-subject PLV correlations, comparison is between Dry vs Gel and Wet vs Gel, in all five frequency bands (Delta, Theta, Alpha, Beta and Gamma).

## Discussion

### Results and Implications

Traditional gel-based systems, while providing high-quality signals, cannot be widely applied to development of EEG outside the lab, notably for home BCI use, due to their complexity, setup time, and maintenance requirements. Dry electrode systems offer convenience but often fall short in signal fidelity, limiting their utility in real-world applications. Thus, researchers increasingly seek to compare different EEG devices in order to identify the most effective approach for reliable EEG recording (Ng et al., 2022; Radüntz & Meffert, 2019). Our results suggest that a minimal, low-cost intervention—wetting the hair with water—makes PSD and FC measures from dry electrodes more similar to those obtained with gel electrodes. Hair-wetting also reduces electrode impedance and the proportion of unusable channels. However, hair-wetting does not yet achieve the signal quality of traditional gel-based EEG systems. These improvements occur despite impedance remaining higher than with gel electrodes and other limitations of dry electrode recording, suggesting that hair-wetting may improve data quality when gel-based recording is impractical.

These findings have important implications for the future of accessible neurotechnology. As interest grows in home-based brain monitoring and BCI applications, particularly for neurofeedback and mental health interventions, the need for practical and reliable EEG acquisition methods becomes increasingly pressing (Nicolas-Alonso & Gomez-Gil, 2012). Improving the balance between usability and signal quality is therefore particularly relevant for scalable EEG applications outside conventional laboratory and clinical settings. Such improvements may facilitate the development of more robust and autonomous neurofeedback solutions that can be deployed outside clinical settings (Biondi et al., 2024).

### Methodological Considerations

While the current results demonstrate the feasibility of the proposed hair-wetting approach, several limitations should nevertheless be considered. First, the recordings were acquired in a fixed dry–wet–gel order in order to minimize potential carry-over between preparation methods, particularly the influence of hair wetting and subsequent gel application across conditions. Although this ordering was deliberately chosen to reduce cross-condition contamination, it also means that condition effects cannot be completely disentangled from potential time or order effects. Second, the dry/wet and gel conditions were recorded using different electrode systems, meaning that hardware-specific characteristics may have contributed to differences with the gel reference despite restricting the analysis to common electrodes. Third, the study was limited to relatively short resting-state recordings performed within a single controlled session. It therefore remains to be determined whether the benefits of hair wetting persist during longer recordings, active cognitive tasks, movement, or repeated sessions (Ehrhardt et al., 2024). Hair characteristics such as length, density, and texture were also not systematically quantified and may influence the effectiveness of hair wetting by affecting electrode–scalp contact (Fiedler et al., 2018). Finally, the availability of auxiliary physiological signals such as EOG and ECG would likely further enhance artifact correction, as supported by the HAPPE framework (Gabard-Durnam et al., 2018). In the present study, however, these signals were intentionally not recorded in order to approximate a realistic and user-friendly home-based recording scenario, where the use of additional sensors over extended periods may be impractical.

### Conclusions and future directions

This study demonstrates that hair-wetting prior to administering dry electrodes substantially improves EEG signal quality, reducing electrode impedance and unusable channels while yielding spectral and functional connectivity measures that more closely resemble a conventional gel-based reference. Although further validation in user-friendly and longitudinal home-based studies is needed, hair wetting represents a simple and practical strategy for improving dry-electrode EEG acquisition in real-world applications. Future developments may also benefit from integrating acquisition-side improvements with lightweight online denoising approaches. For example, the generalized eigen decomposition based pipeline (Ros et al., 2025) enables rapid and robust EEG denoising and could potentially complement the present acquisition-side approach in real-time neurofeedback and BCI applications.

**Table S1.**
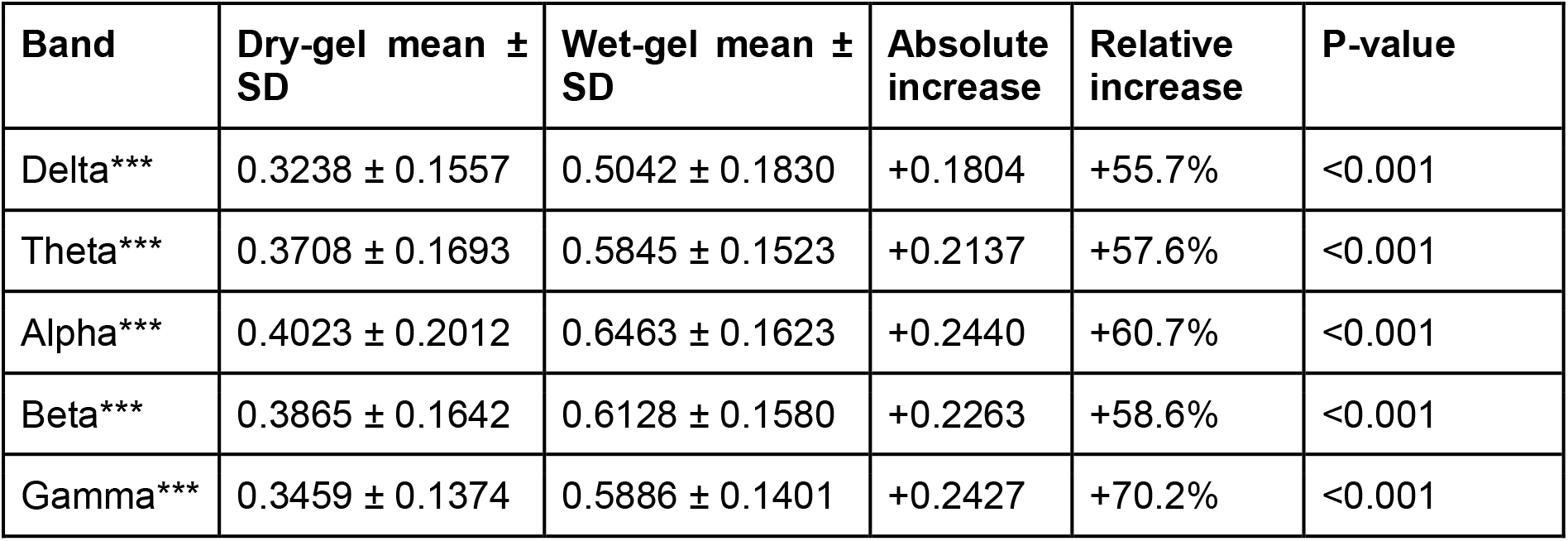
Mean functional connectivity correlations with the gel condition and relative improvement after hair wetting across EEG frequency bands.

